# A low-threshold potassium current enhances sparseness and reliability in a model of avian auditory cortex

**DOI:** 10.1101/249011

**Authors:** Margot C. Bjoring, C. Daniel Meliza

## Abstract

Birdsong is a complex vocal communication signal, and like humans, birds need to discriminate between similar sequences of sound with different meanings. The caudal mesopallium (CM) is a cortical-level auditory area implicated in song discrimination. CM neurons respond sparsely to conspecific song and are tolerant of production variability. Intracellular recordings in CM have identified a diversity of intrinsic membrane dynamics, which could contribute to the emergence of these higher-order functional properties. We investigated this hypothesis using a novel linear-dynamical cascade model that incorporated detailed biophysical dynamics to simulate auditory responses to birdsong. Neuron models that included a low-threshold potassium current present in a subset of CM neurons showed increased selectivity and coding efficiency relative to models without this current. These results demonstrate the impact of intrinsic dynamics on sensory coding and the importance of including the biophysical characteristics of neural populations in simulation studies.

## Introduction

Decoding a speech stream requires the auditory system to recognize phonemes—invariant auditory objects that do not change their identity even when different contexts or speakers cause shifts in their underlying acoustic structure ***(Liberman et al., 1961; Iverson and Kuhl, 2000)***. In animal models, higher-order auditory areas show increased selectivity and tolerance ***(Tsunada et al., 2012; Meliza and Margoliash, 2012)***, which could facilitate such invariant auditory object recognition. Selectivity, or sparseness, is a metric that indicates when a neuron responds strongly only to a small subset of stimuli ***(Rolls and Tovee, 1995; Vinje and Gallant, 2000)***, which could facilitate discriminations between complex vocalizations. Tolerance is a complementary metric that indicates how invariant a neuron’s response is to variations in the stimulus that do not affect identity ***(Zoccolan et al., 2007)***. Despite the theoretical importance of selectivity and tolerance to high-level sensory coding ***(Riesenhuber and Poggio, 1999)***, the mechanism of their emergence in the processing hierarchy remains unknown.

Songbirds communicate with acoustically complex vocalizations, which requires them to perform many of the same kinds of auditory discrimination tasks as humans ***(Gentner, 2004)***. In the avian auditory system, the caudal mesopallium (CM) is a cortical-level area that contains a population of neurons highly selective for particular song elements, yet tolerant of low-level acoustic differences between renditions ***(Gentner and Margoliash, 2003; Meliza and Margoliash, 2012)***. In contrast, the neurons in Field L, immediately upstream of CM, show low selectivity and tolerance ***(Calabrese and Woolley, 2015)***. In the classical model of selectivity, complex feature representations emerge through feedforward synaptic connections that pool sparsely from upstream sources ***(Hubel and Wiesel, 1965)***. This model does not match experimental evidence from CM, however, which shows that selective neurons receive a more distributed pool of inputs than the sparse model would predict ***(Perks and Gentner, 2015)***. A distributed scheme of selectivity could arise from nonlinear dynamics within the neurons themselves, but the mechanisms of this have not been explored.

In many of the auditory areas in the hindbrain and midbrain, nonlinear neural dynamics profoundly affect how a variety of low-level acoustic features are encoded ***(Carr and Soares, 2002; Khurana et al., 2011; Gai et al., 2014)***. For example, neurons that express a low-threshold potassium current (*I_K_LT__*) produce highly phasic responses, responding to rapid increases in excitation but not to slow or steady-state depolarizations ***(Rothman and Manis, 2003; Gai et al., 2009)***. These dynamics are critical to temporal precision in sound localization circuits ***(Carr and MacLeod, 2010; Gai et al., 2014)*** and can enhance signal detection in noisy conditions ***(Svirskis et al., 2002)***. Could intrinsic dynamics also contribute to sensory processing at cortical levels? In CM, the putatively excitatory neurons exhibit diverse intrinsic firing patterns ***(Chen and Meliza, 2017)***. About 30% exclusively produce phasic responses that depend on a low-threshold potassium current, whereas the remainder produce mostly tonic responses. The functional significance of this diversity remains unclear.

The goal of the present study is to understand how *I_K_LT__* and the phasic dynamics it produces could contribute to the emergence of selectivity and tolerance in CM. To investigate how intrinsic dynamics could interact with sensory-driven inputs, we developed an auditory response model that combines a spectrotemporal receptive field (RF) with a biophysical spike generation mechanism. In this linear-dynamical cascade model, the RF component approximates the integration over multiple spectral channels and time lags performed by the circuits upstream of the neuron and by the neuron’s dendritic tree. This part of the model is identical to the linear filter component of the classic linear-nonlinear Poisson (LNP) model ***(Theunissen et al., 2000; Pillow et al., 2005)***. However, instead of using the output of the filter as the conditional intensity of a probabilistic spiking process, we feed it into a single-compartment, biophysical model that produces spikes through Hodgkin-Huxley dynamics. This approach allows us to integrate our knowledge about intracellular properties in CM into a model for high-level coding properties.

## Results

***Figure 1***A represents the operation of the linear-dynamical cascade model. The spectrogram of a zebra finch song stimulus was convolved with the RF to produce a driving current ***I***_*stim*_(*t*), which was injected into a single-compartment dynamical neuron model containing several voltage-dependent sodium and potassium currents. The mathematical descriptions of these currents were derived from an intracellular study of the excitatory neurons in the caudal mesopallium (CM) of zebra finches ***(Chen and Meliza, 2017)***. A key feature of this model is that it can produce phasic or tonic responses depending on the conductance of a low-threshold potassium current (*g_K_LT__*), which is experimentally known to be present in some CM neurons.

**Figure 1.**
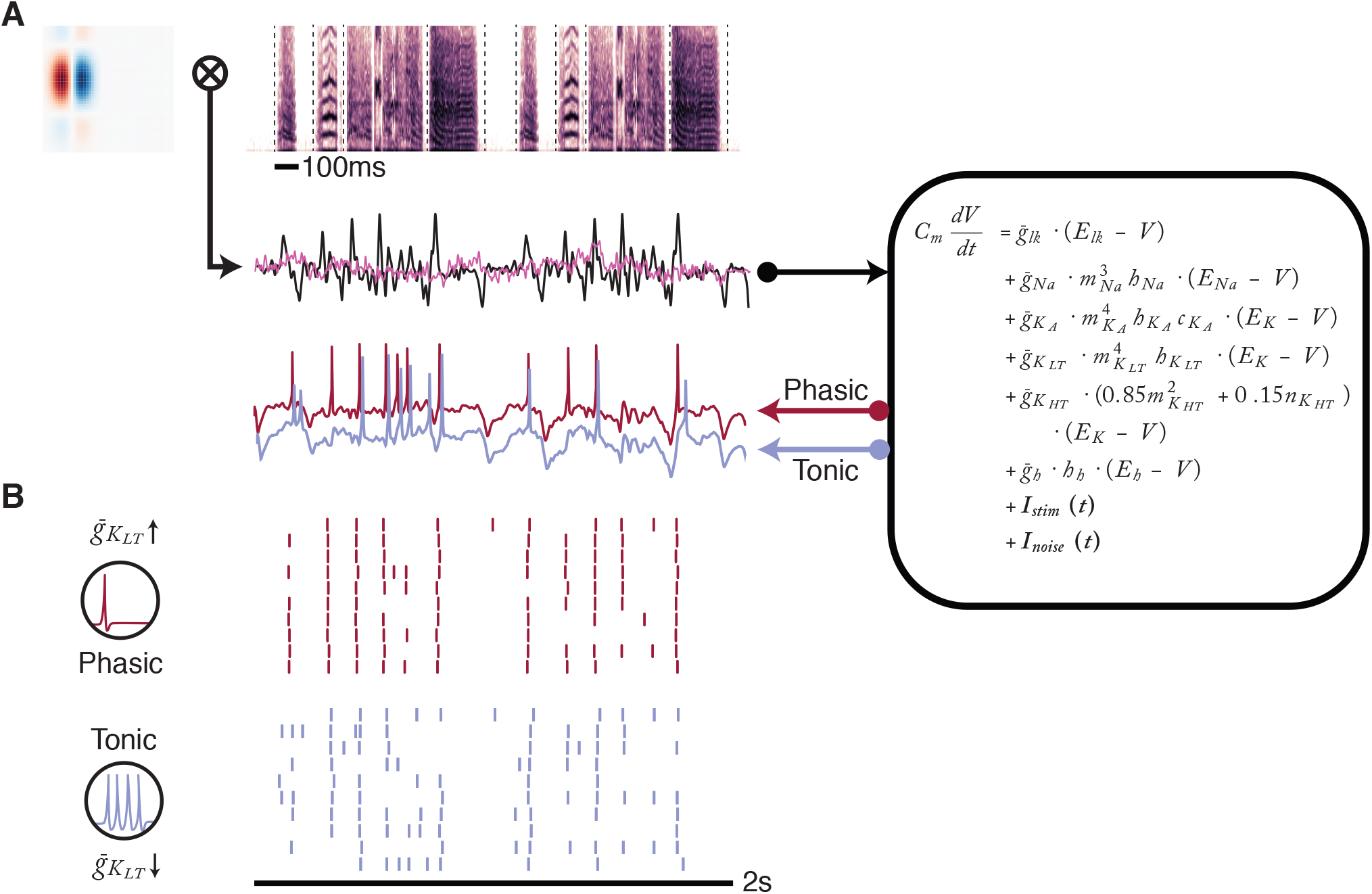
Simulating auditory responses with biophysical dynamics. A) Auditory responses were simulated by convolving a spectrotemporal receptive field (upper left) with the spectrogram (upper right) of an auditory stimulus, in this case a zebra finch song. Black dashed lines indicate syllable boundaries. The resulting convolution (black line) provided the driving current (***I***_*stim*_(*t*)) to the single-compartment biophysical model used in this study (right). Low-pass-filtered pink noise (pink line) was added in each trial as a stimulus-independent current (***I***_*noise*_(*t*)). The model was integrated to produce a simulated voltage trace (lower left). The conductance of a low-threshold potassium channel parameter (***g_K_LT__***) was set to 0 nS for tonic dynamics (blue line) or 100 nS for phasic dynamics (red line). Voltage traces are vertically offset for better visibility. B) Raster plots of the full simulation for the stimulus-RF pair in A across 10 trials for phasic (red) and tonic (blue) model. The example demonstrates how the phasic model produces more reliable and temporally precise responses than the tonic model.

As seen in ***Figure 1***B, the model’s intrinsic dynamics affected its response to zebra finch song. When ***g_K__LT_*** was high (phasic model), responses were more precise and more reliable than when ***g_K_LT__*** was absent (tonic model). This effect was consistent across RFs with different spectral and temporal parameters and across multiple stimuli (***Figure 2***). It is important to note that although ***I_K_LT__*** is an outward current with an overall hyperpolarizing influence, increasing it did not simply reduce excitability. In some cases, the phasic model responded at a much lower rate and to only a handful of notes across the stimulus set (***Figure 2***B). In other cases, the firing rates of the phasic and tonic models were nearly the same (***Figure 2***A,C), and the effects of manipulating were primarily on trial-to-trial variability. Overall, the average difference in firing rate between tonic and phasic models was 0.90 ± 1.06 Hz (*t*_59_ = 6.58; *p* < 0.001).

**Figure 2.**
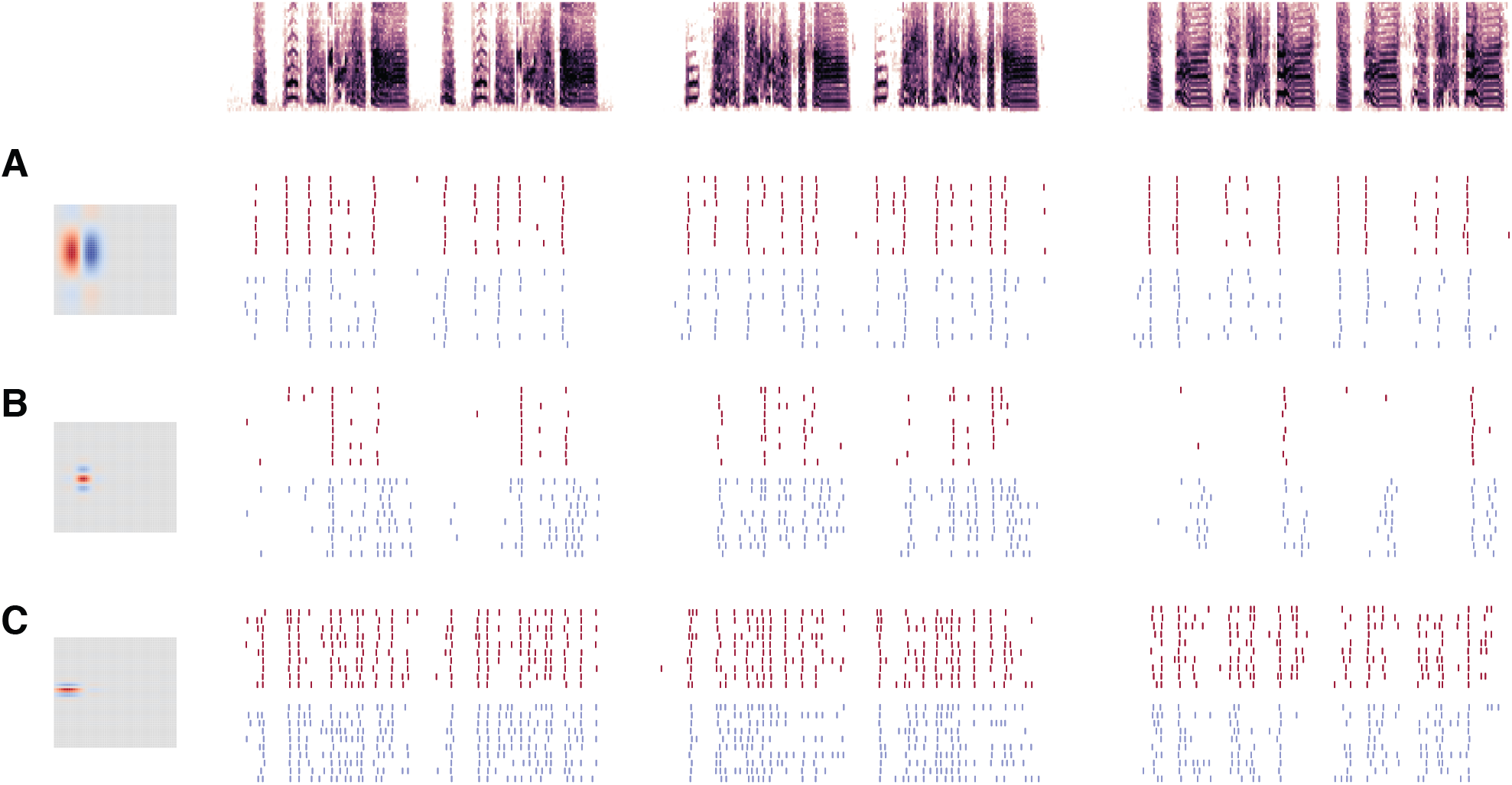
Responses of simulated neurons. Raster plots for models with three different RFs (rows, A-C) in response to three different stimuli (columns). As in Fig. 1, raster plots show 10 simulated responses of models with phasic (red) or tonic (blue) dynamics. Different RFs produce different levels of sparseness, selectivity, and precision in the spiking responses, but tonic models are consistently less precise and less selective than phasic models.

To investigate how phasic excitability affects functional response properties, in this study we focused on rate-based theories of sensory coding. The fundamental idea of rate coding is that neurons convey information about sensory inputs by modulating their average firing rate over relatively long intervals. Following previous studies ***Jeanne et al., 2011; Meliza and Margoliash, 2012)***, we defined these intervals by dividing the responses into segments corresponding to song syllables (as in ***Figure 1***), which are well-defined units of zebra finch song that convey information about individual identity ***(Cynx, 1990)***. We then calculated metrics based on how the average rates within those intervals were distributed across syllables and trials (***Figure 3***A).

**Figure 3.**
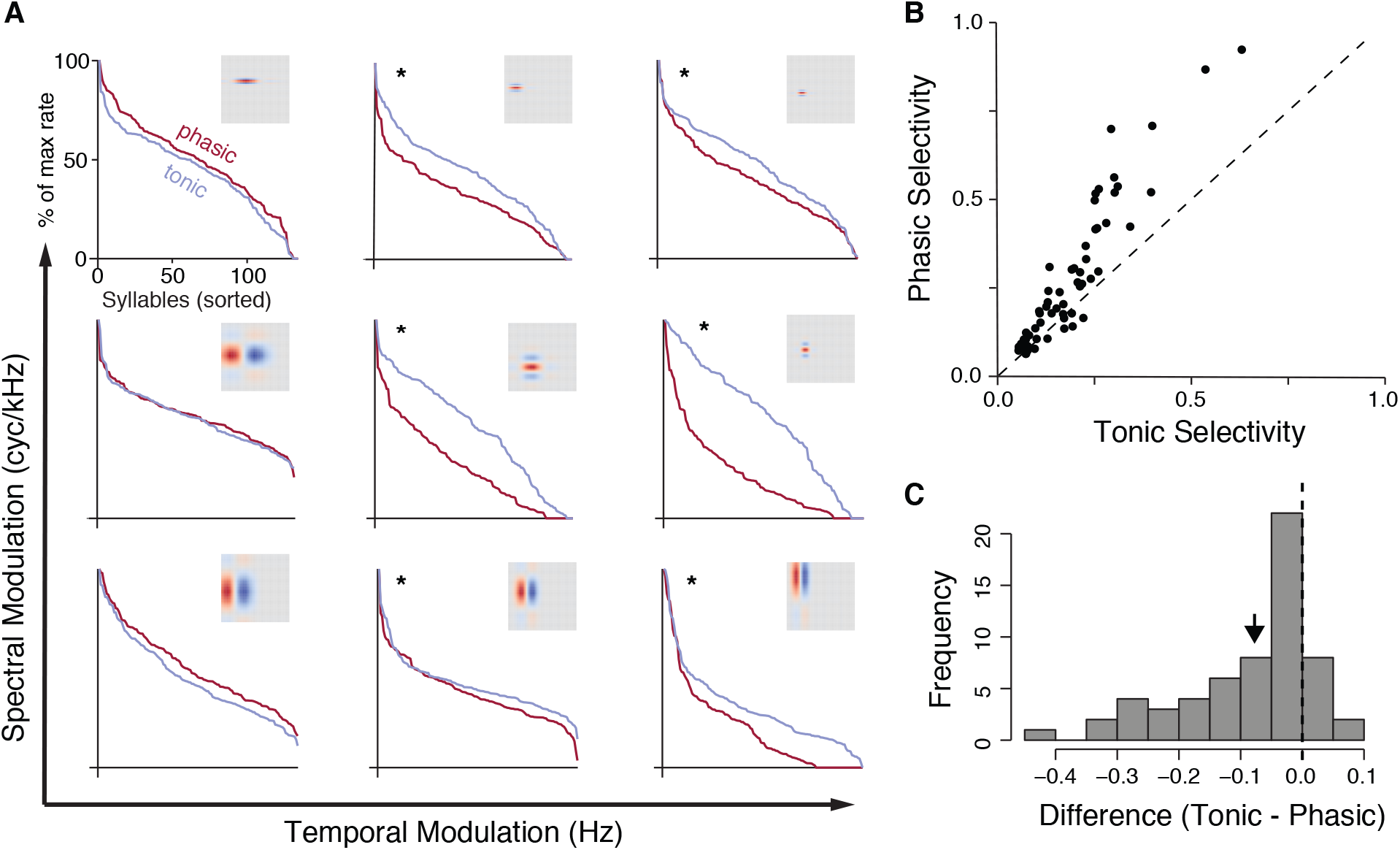
Phasic dynamics increase selectivity. A) Normalized cumulative distribution of response rates for nine example RFs with phasic (red) and tonic (blue) dynamics. High selectivity is indicated by a response distribution with a heavy tail on the left side of the plot, indicating that only a few stimuli account for the strongest responses. Asterisks indicate a significant difference in the distribution of response rates (*p* < 0.05, Komolgorov-Smirnov test). Receptive fields are arranged by increasing spectral and temporal modulation frequency. B) Selectivity of paired phasic and tonic models. Selectivity is quantified using activity fraction (see Methods). Each point corresponds to one RF. The dashed reference line indicates the line of equal selectivity. Models with phasic dynamics are consistently more selective than their tonic counterparts (*t*_59_ = 6.47; *p* < 0.001). C) Histogram of the difference in selectivity between phasic and tonic models. Negative values indicate the phasic model has higher selectivity than the tonic model. The black arrow shows the mean difference.

We first examined whether phasic dynamics made neurons more selective. Selectivity, also known as lifetime sparseness, is a well-established rate-coding metric defined as the tendency of a neuron to respond strongly to only a small subset of stimuli ***(Rolls and Tovee, 1995)***. A selective neuron has a skewed, heavy-tailed response distribution in which a few stimuli account for the strongest responses, and the remainder produce only weak excitation. In contrast, a less selective neuron has a small-tailed, Gaussian response distribution, with most of the responses concentrated around the mean. For the examples shown in ***Figure 3***A, the models with phasic dynamics tended to be significantly more selective than the corresponding models with tonic dynamics. This effect was not uniform, but appeared to be stronger in the models with higher temporal-modulation frequencies in their RFs.

To draw more general conclusions, we simulated responses using models comprising 60 different RFs that matched the distribution of spectral and temporal parameters seen in the zebra finch auditory cortex ***(Woolley et al., 2009)***. Each RF was combined with phasic and tonic dynamics for a total of 120 models. Across this population, phasic models were consistently more selective than tonic models with the same RFs (***Figure 3***B and C).

We also observed that phasic models were more reliable across trials, indicating that they were less affected by the noise current ***I***_*noise*_. To quantify this effect in the context of rate coding, we calculated the mutual information (MI) between the response rate and syllable identity. MI is defined as the difference between the response (total) entropy, which represents how much information the neuron can carry based on its range of firing rates, and noise (or conditional) entropy, which represents how much information is lost due to the variability of a neuron’s response to the same stimulus. A neuron that responds more similarly across trials will have a low noise entropy, bringing its MI closer to its total entropy.

***Figure 4*** illustrates how intrinsic dynamics affected the distribution of response rates across trials for the same nine example RFs shown previously. The tonic models tended to have a higher total entropy across these examples, although in several cases, the phasic models are slightly higher. However, for all of these examples, the response rates of the phasic models cluster more closely around the mean response, resulting in lower noise entropy compared to the tonic models.

**Figure 4.**
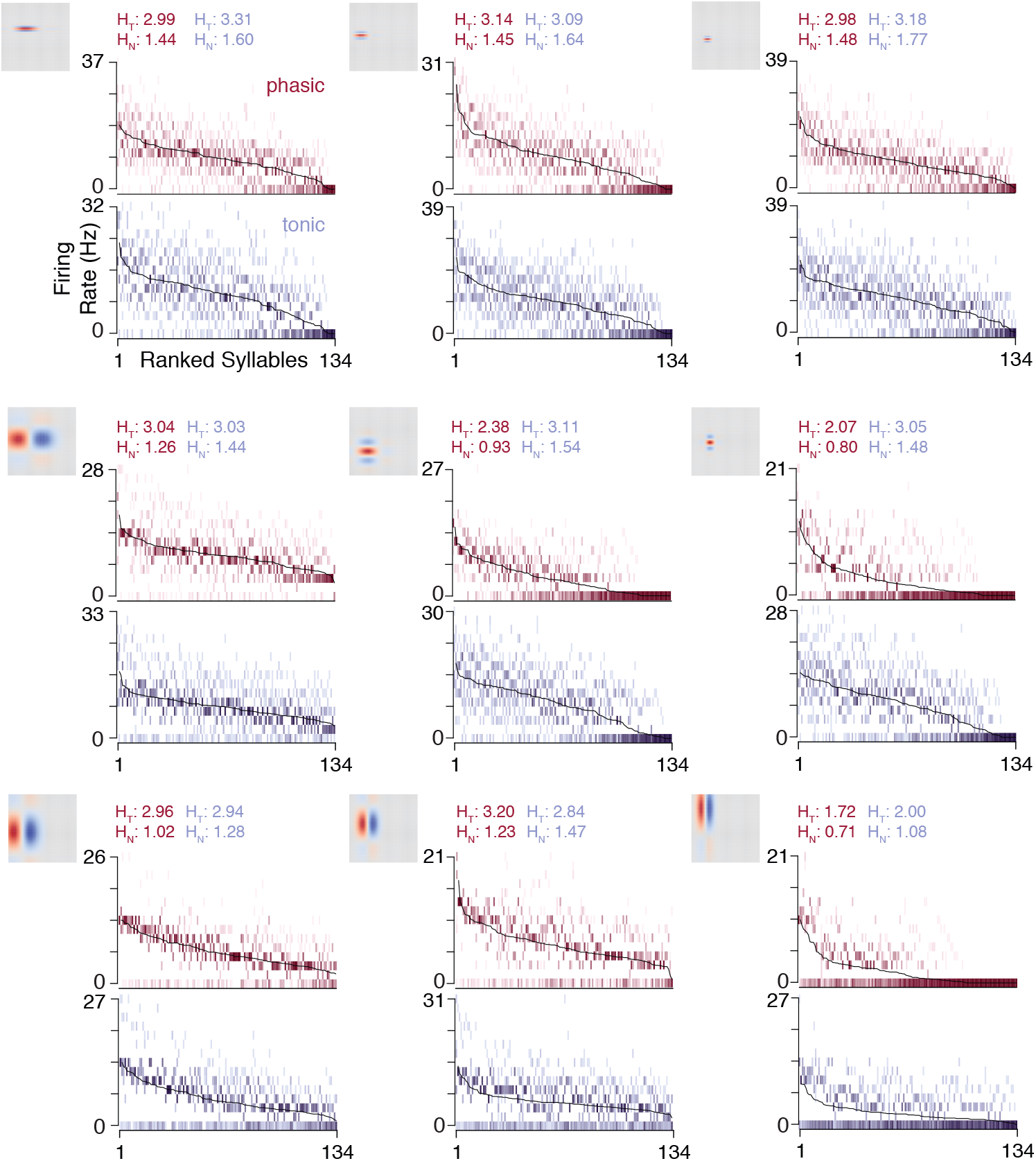
Phasic dynamics reduce trial-to-trial variability in spike rates. Full response distributions for the nine example neurons shown previously. Response rates are calculated for each syllable and trial and discretized into 15 bins. The black lines indicates the average across trials; the spread of response rate bins around those lines show the trial-to-trial variability of the response rates. Phasic models tended to show less variability around the mean rate than tonic models. Total entropy (***H***_*T*_) and noise entropy (***H***_*N*_) values are shown above each plot (phasic: red; tonic: blue).

Across all 60 RF pairs, the total entropy was slightly higher for tonic neurons (***Figure 5***A,B). That advantage was more than canceled out by a much larger increase in noise entropy(***Figure 5***C and D). Every phasic simulation had lower noise entropy than its tonic pair. The net effect was that phasic models had higher MI than the corresponding tonic models (***Figure 5***E, F).

**Figure 5.**
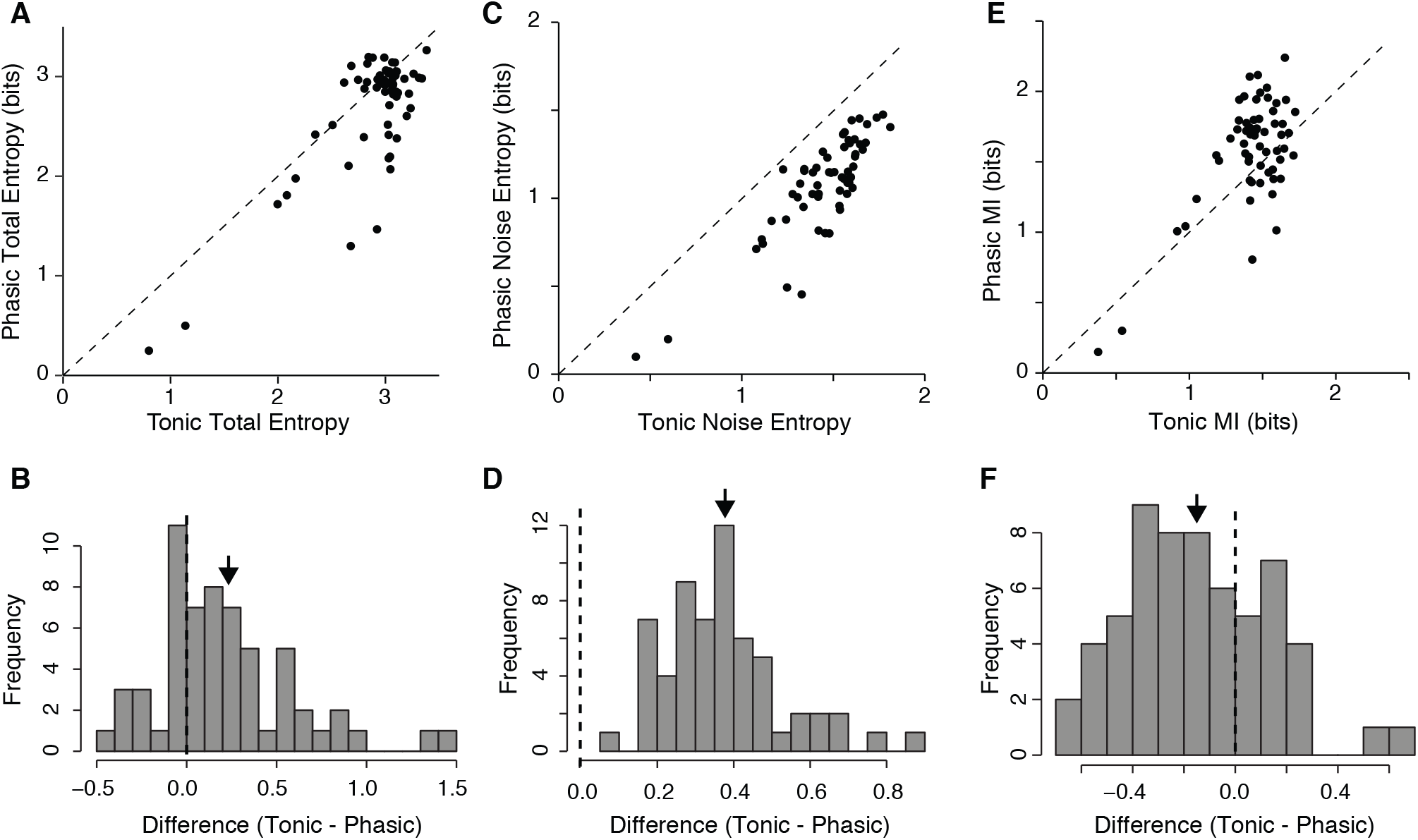
Phasic dynamics confer increased mutual information about syllable identity by increasing reliability. A) Comparison of total (response) entropy, which represents the maximum information capacity of the model. Phasic and tonic models had similar total entropy, though tonic models had a slight advantage (*t*_59_ = 4.41;*p* < 0.001). B) Histogram of the difference in total entropy between phasic and tonic models. Positive values indicate the tonic model had higher total entropy than the phasic model. C) Comparison of noise entropy, which represents variation in responses to the same stimulus across trials. Noise entropy decreases the amount of information conveyed from the theoretical maximum. All of the phasic models had lower noise entropy than tonic models (*t*_59_ = 18.79;*p* < 0.001). D) Histogram of the difference in noise entropy between phasic and tonic models. E) Mutual information of paired phasic and tonic models. On average, models with phasic dynamics had higher mutual information than the corresponding tonic models (*t*_59_ = −4.21;*p* < 0.001). F) Histogram of the difference in mutual information between phasic and tonic models.

To ensure that these results were not peculiar to the signal-to-noise ratio (SNR) chosen for this simulation, we repeated these analyses with simulations of different SNR values (***Figure 6***A-D). Phasic models consistently had lower noise entropy and selectivity, and only at the highest and lowest SNR values did phasic and tonic MI approach each other.

**Figure 6.**
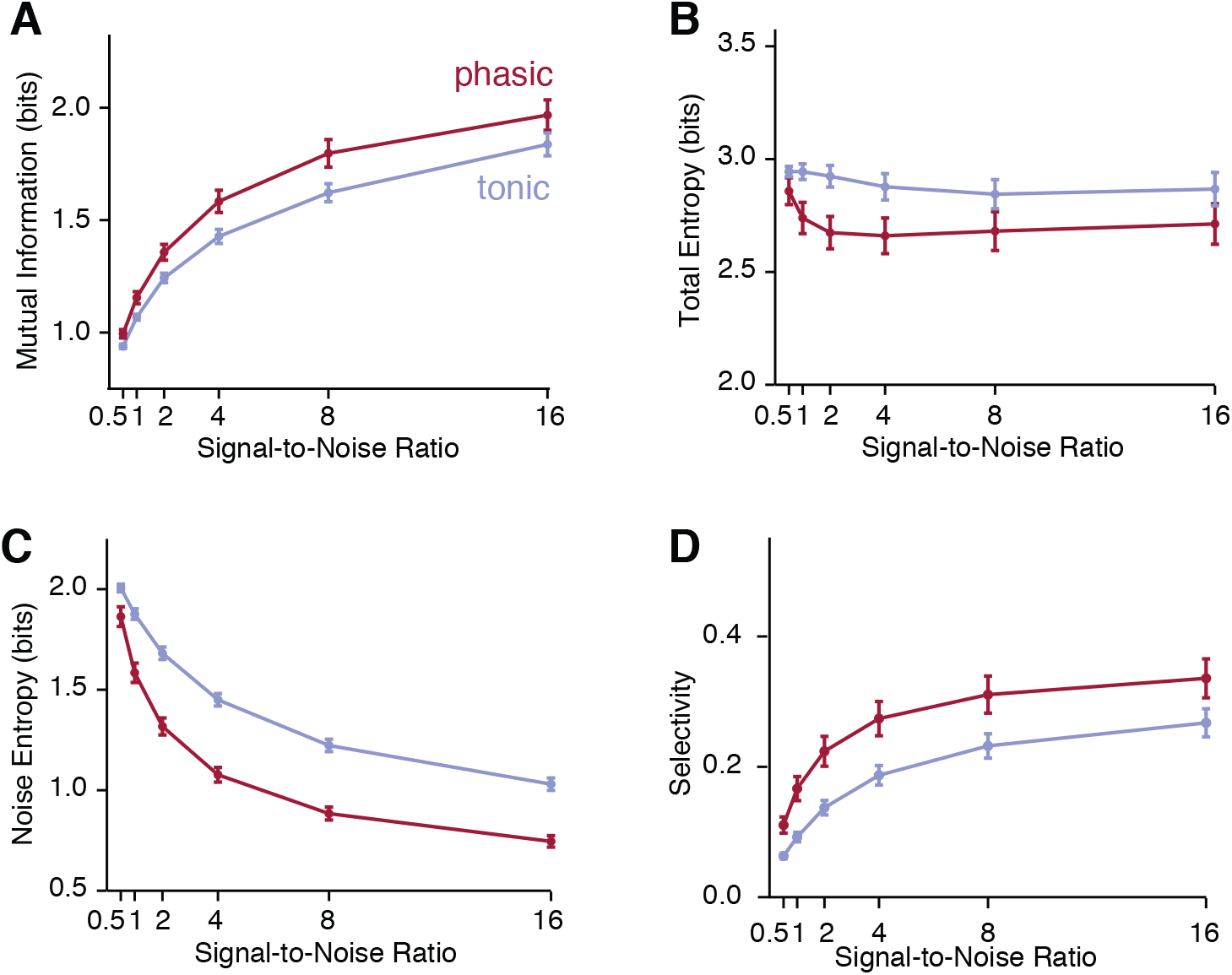
Signal-to-noise ratio does not change the relationship between phasic and tonic models. A) The effect of different signal-to-noise ratios (SNR) on the outcome of the mutual information analysis. As the amplitude of the noise current increases relative to the amplitude of the stimulus-driven current (decreasing SNR), mutual information decreases, but phasic models continue to outperform tonic models. The red line shows the mean mutual information for phasic models, and the blue line shows the mean for tonic models. The bars show standard error. B) The effect of SNR on total entropy. The red line shows the mean total entropy for phasic models, and the blue line shows the mean for tonic models. The bars show standard error. C) The effect of SNR on noise entropy. The red line shows the mean noise entropy for phasic models, and the blue line shows the mean for tonic models. The bars show standard error. D) The effect of SNR on selectivity for phasic (red) and tonic (blue) models. The line shows mean selectivity and the bars show standard error.

How are selectivity and MI related? In these simulations, total entropy and selectivity were negatively correlated (***Figure 7***A). This effect is unsurprising given that selective neurons respond similarly to a large proportion of stimuli. They encode more information about a few stimuli at the expense of encoding less information about the entire stimulus set. However, when we consider coding efficiency, which quantifies how much of this potential bandwidth is actually used instead of being lost to noise (***Figure 7***C), two trends emerge. First, tonic models have lower coding efficiency than phasic models, consistent with the observation that they have higher noise entropy and lower MI. Second, only phasic neurons are able to achieve both high selectivity and high coding efficiency.

**Figure 7.**
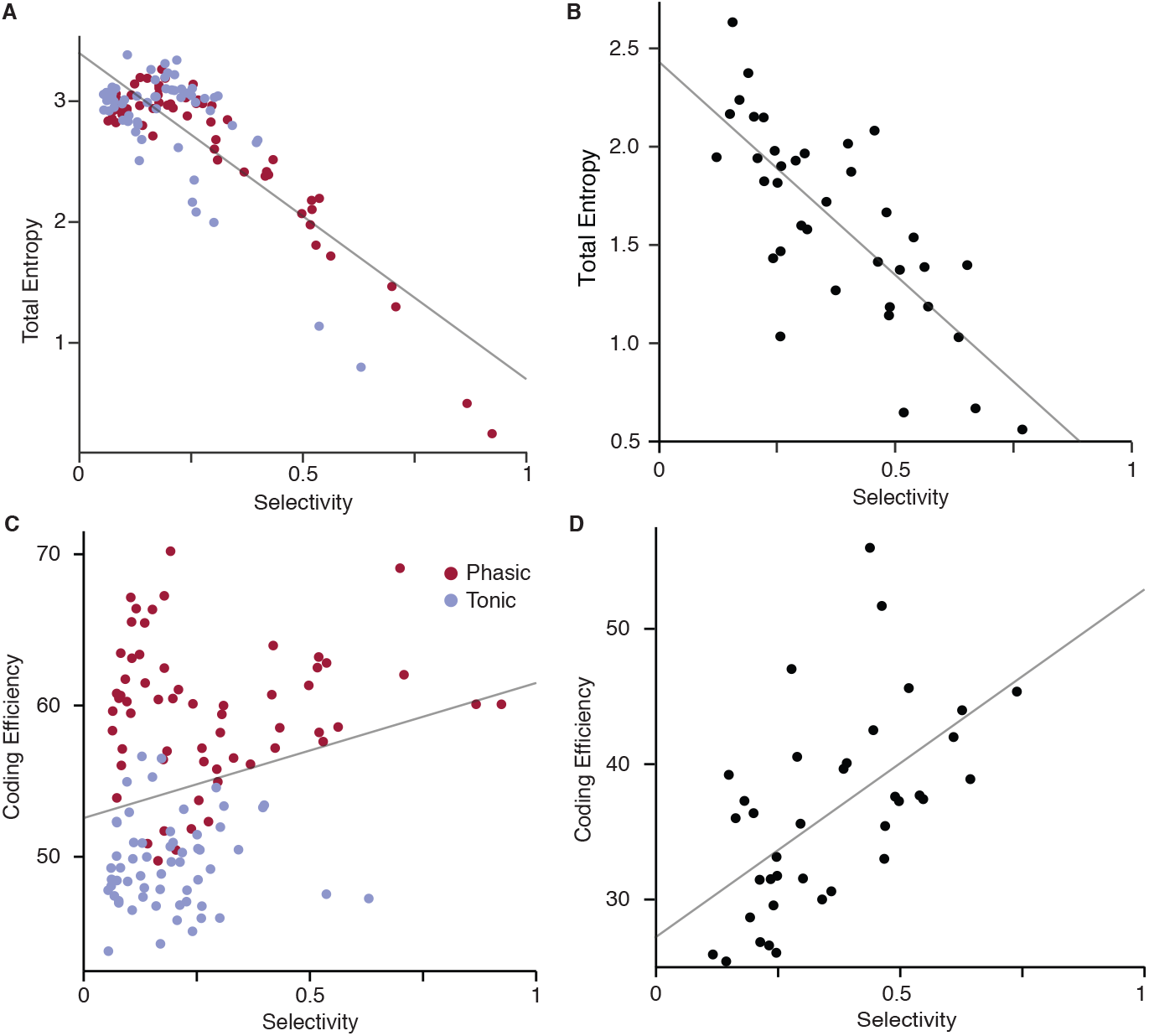
The model predicts a relationship between coding efficiency and selectivity. A) Total entropy and selectivity are negatively correlated for both phasic and tonic models, showing the inherent trade-off between selectivity and entropy (Pearson *r* = -0.84; *p* < 0.001). B) CM neurons in zebra finches exhibit the same trade-off (Pearson *r* = -0.76; *p* < 0.001). Data from ***Theunissen et al. (2011)***. C) Coding efficiency, calculated as the percentage of total entropy not lost to noise, is enhanced by phasic dynamics. Tonic models (blue points) cluster around lower values of coding efficiency and also lack the population of high selectivity neurons that the phasic models (red points) exhibit. The population of models show a weak positive correlation between coding efficiency and selectivity (Pearson *r* = 0.24; *p* = 0.008). D) CM neurons in zebra finches show a stronger positive correlation between selectivity and coding efficiency (Pearson *r* = 0.58; *p* < 0.001), as they lack the population of low-selectivity, high-reliability neurons seen in the model (C). Data from ***Theunissen et al. (2011)***.

As an independent test of the validity of this model, we applied the same selectivity and MI analyses to a public corpus of recordings from zebra finch CM ***(Theunissen et al., 2011)***. The relationships between selectivity and MI predicted by the model were largely borne out in the experimental data. We observed a similar tradeoff between selectivity and total entropy (***Figure 7***B), and an even stronger positive correlation between selectivity and coding efficiency (***Figure 7***D). The average coding efficiency was lower in the experimental data, which likely reflects additional sources of variability *in vivo*. Selectivity was somewhat higher *in vivo*, perhaps because of additional nonlinearities in the actual CM receptive fields. Interestingly, the cluster of models with high coding efficiency and low selectivity seen in the simulated data is not present in the experimental data.

Why are models with phasic dynamics consistently more reliable and selective than models with tonic dynamics? Low-threshold potassium currents counteract the regenerative sodium current produced during spike initiation. As a result, cells expressing these currents only spike in response to a rapid increase in excitation. These dynamics enable phasic neurons in the auditory hindbrain and midbrain to detect coherent excitation with high temporal precision, even in noisy conditions ***(Rathouz and Trussell, 1998; Svirskis et al., 2002; Khurana et al., 2012)***. Consistent with this idea, we observed in our sensory model that moments of high concordance between the RF and the stimulus created peaks in the driving current ***I***_*stim*_(*t*). The phasic models spiked almost exclusively at these moments (***Figure 8***A). In contrast, the tonic models were more sensitive to the absolute level of ***V***(*t*) and increased their firing rate as the integral of ***I***_*stim*_(*t*)became more positive (***Figure 8***B).

**Figure 8.**
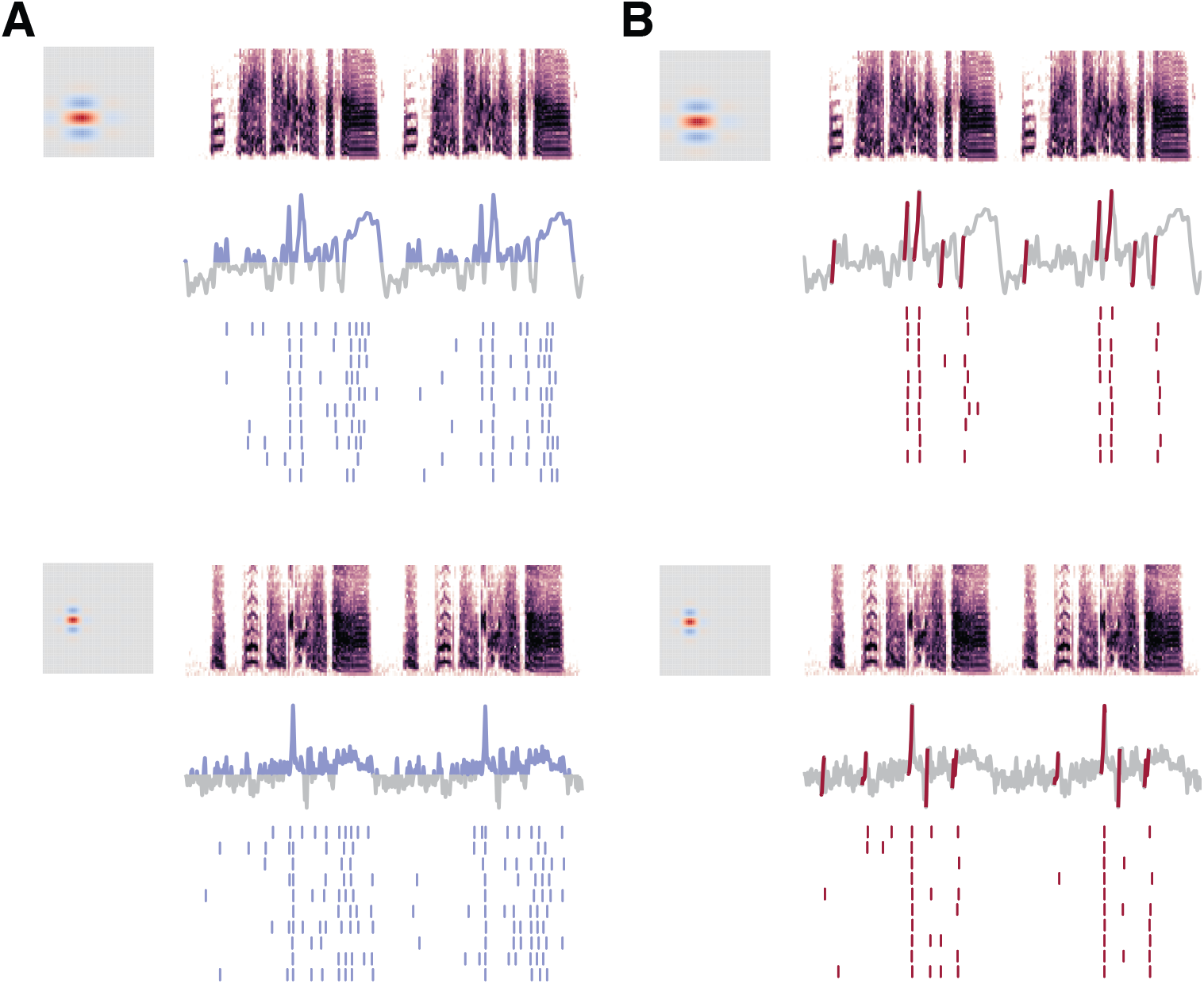
Phasic selectivity is driven by slope detection. A) Tonic models responded to the level of excitation in the driving current. Spiking activity aligned well to the moments when net excitation was greater than zero, and the model produced jittery, unreliable responses to broad peaks of excitation. Blue segments of the trace mark when the convolution is positive. B) Phasic models only responded when driving current contained a positive slope with a high rate of change. Spike times aligned well to these slopes. The red segments of the convolution mark the times at which the difference of the smoothed convolution was 1.5 standard deviations above the mean difference. Because such slopes were relatively scarce and brief, the spikes predicted by the phasic model were selective and had little temporal variation.

Based on this observation, we hypothesized that the effect of phasic excitability on selectivity would be stronger when the RF produced a driving current with sparse peaks of excitation. In the spectral domain, this corresponds to convolutions that have more power at higher frequencies and RFs that have higher temporal modulation frequencies. To test this, we simulated another set of data with eight RFs, holding all RF parameters constant except temporal modulation (Ω_*t*_), which we varied between 10 and 80 Hz.

At low values of Ω_*t*_, convolutions of the RF and a stimulus produced mostly slow modulations, and the response of phasic and tonic models were similar, as the example in ***Figure 9***A shows. As the modulations in the convolution became faster and large deflections became sparser (***Figure 9***A, 80 Hz), the response of the phasic model became more selective, while the tonic model, responding to the shape of the slow modulation still present in the signal, showed no inclination toward selectivity. ***Figure 9***B presents the full set of simulations, showing a strong interaction between the temporal modulation of the RF and selectivity. Phasic models showed a strong increase in selectivity as Ω_*t*_ of the RF increased but tonic models were unaffected.

**Figure 9.**
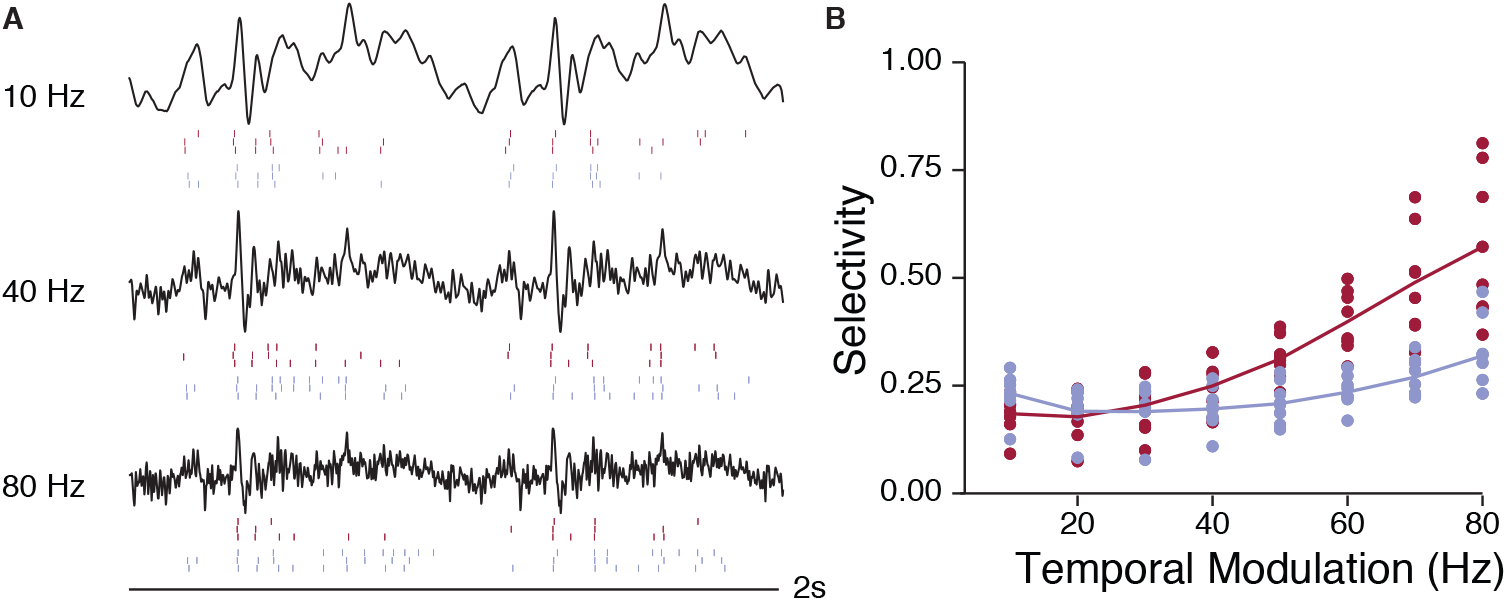
Selectivity depends on an interaction between phasicness and RF shape. A) Example convolutions of an RF with Ω_*t*_ = 10,40, and 80 Hz. Rasters of phasic (red) and tonic (blue) responses for 3 of 10 trials are shown below. At 10 Hz, slow modulations predominate. At 40 Hz, the convolution contains more sharp peaks. At 80 Hz, the peaks become sparser, but only the response of the phasic example becomes more selective. B) The selectivity of neural simulations increases as the temporal modulation of the RF (Ω_*t*_) increases, but only for phasic models. The selectivity of tonic simulations shows little effect. The interaction between model type and Ω_*t*_ is significant (*F*_1,124_ = 54; *p* < 0.001)

## Discussion

This study investigated how intrinsic dynamics could interact with spontaneous and sensory-driven synaptic currents in CM, a cortical-level auditory area. We used a novel linear-dynamical cascade model that combines a spectrotemporal receptive field with a biophysical description of intrinsic membrane dynamics. We found that a low-threshold potassium current (***I_K_LT__***) expressed in a subset of CM neurons ***(Chen and Meliza, 2017)*** strongly affected how the model neurons encoded information about song stimuli in firing rates. Phasic models (with high ***I_K_LT__***) were more selective and more tolerant of noise than models with identical RFs and tonic dynamics. Furthermore, a population of phasic and tonic models reproduced the distribution of selectivity and coding efficiency seen *in vivo* (***Figure 7***). These results suggest that a diversity of intrinsic intracellular properties contributes to the emergence of higher-order functional properties in CM.

The model we developed for this analysis is a special case of the linear-nonlinear (LN) cascade model used in many studies of sensory coding (e.g., ***Eggermont et al., 1983; Keat et al., 2001; Schwartz et al., 2006***). In the standard LN model, the nonlinearity is a history-independent function that transforms the output of the linear stage into an instantaneous spiking probability. More recent LN models incorporate history dependence through a linear kernel convolved with past spike times, as in the generalized linear model ***(Paninski, 2004; Pillow et al., 2005)***, or through non-biological, dynamical state variables, as in the spike response and generalized integrate-and-fire models ***(Įolivet et al., 2004; Kobayashi et al., 2009; Lynch and Houghton, 2015)***. The present study required a more biologically realistic representation of the dynamics so that we could manipulate a specif c current of interest, ***I_K_LT__***. We therefore used a conductance-based biophysical model to generate spikes. This model lacks many of the morphological and physiological properties of CM neurons and omits any circuit-level organization, but it is capable of reproducing the tonic and phasic firing patterns observed in slices ***(Chen and Meliza, 2017)***. For the linear stage, we convolved the stimulus spectrogram with simplified STRFs from a previous study of Field L ***(Woolley et al., 2009)***. This parametric model of STRF structure allowed us to investigate how different RF features interacted with nonlinear dynamics. Despite its simplicity, the model was able to reproduce qualitatively realistic responses to song stimuli, which strengthens confidence in the biological validity of the effects described here.

The function of ***I_K_LT__*** in shaping auditory responses has been studied extensively in the hindbrain and midbrain ***(Locke and Nerbonne, 1997; Carr and Soares, 2002; Rothman and Manis, 2003; Sivaramakrishnan and Oliver, 2001)***. Because this current rapidly activates at voltages around or below the spike-initiation threshold, it counterbalances low-frequency fluctuations in driving current and makes neurons sensitive to the change in a stimulus rather than its steady-state value ***(Meng et al., 2012)***. When incorporated into our cortical-level sensory model and tested on song stimuli, ***I_K_LT__*** causes responses to become sparser, more precise, and more reliable (***Figure 2***). We focused here on quantifying these effects using rate-based encoding metrics, which are well established in the general literature and for the avian auditory system ***Jeanne et al., 2011; Meliza and Margoliash, 2012)***, though the effects on temporal precision are worthy of future consideration.

Selectivity, which is also called activity fraction or lifetime sparseness ***(Rolls and Tovee, 1995; Vinje and Gallant, 2000)***, was strongly enhanced by ***I_K_LT__*** (***Figure 3***). Syllables that produced strong responses remained strong, but weak responses became weaker, causing response rate distributions to become more heavy-tailed. This led to a reduction in the total entropy of the distributions (***Figure 5***), which reflects the fact that a more selective neuron responds similarly to all the weak stimuli and therefore encodes little information about the majority of the stimulus set. At the same time, the phasic models lost much less of their bandwidth to noise, which led to a net increase in mutual information and coding efficiency. Thus, ***I_K_LT__*** can increase both the sparseness of the neural code and its reliability.

The effects of ***I_K_LT__*** on neural coding were not simply a product of reducing excitability. Rather, they arose from complex interactions between the RF, the stimulus, and the nonlinear dynamics of the membrane. Phasic dynamics did not increase selectivity and coding efficiency uniformly, but instead had stronger effects for RFs that produced convolutions with brief peaks of excitation. These peaks, of course, are what most reliably drive phasic neurons ***(Golding et al., 1995; Meng et al., 2012)***. In this study, the RFs most strongly affected by dynamics had bandpass temporal characteristics. Zebra finch song has strong, broadband rhythmic structure, and bandpass RFs produce sharp peaks at syllable onsets and offsets ***(Woolley et al., 2009)***. We expect that had we been able to include more complex, multi-feature RFs like those recently reported in starling CM and NCM ***(Kozlov and Gentner, 2016)***, we would have seen an even stronger effect of dynamics on the brief moments when the RFs matched the stimulus.

Neurons in the avian auditory pallium exhibit a broad range of functional response properties. Even within increasingly higher-order areas, some neurons are strongly selective and others are not ***(Gentner and Margoliash, 2003; Meliza et al., 2010; Meliza and Margoliash, 2012)***. The cause of this diversity, which is also seen in other secondary areas ***(Cohen, 2004; Zoccolan et al., 2007)***, is unclear. The models in this study sampled from a distribution of RFs with a broad range of spectral and temporal statistics, but only when the population included both phasic and tonic dynamics did we observe the broad, correlated distribution of selectivity and coding efficiency seen in zebra finch CM (***Figure 7***). This result supports the idea that intrinsic dynamics may contribute to functional diversity ***(Padmanabhan and Urban, 2010)***, and may help to explain why CM contains both tonic and phasic neurons. Interestingly, the model response properties were more diverse than seen *in vivo*, suggesting that intrinsic dynamics may be somehow matched to RF properties.

Animals from a broad range of species are able to perform invariant object recognition in challenging conditions ***(Logothetis and Sheinberg, 1996; Griffiths and Warren, 2004)***. In many sensory pathways, there is a hierarchical increase in selectivity and tolerance for natural objects that is thought to underlie this remarkable ability. However, the circuit-level implementation of this computation remains largely theoretical ***(DiCarlo et al., 2012)***. Although many models have followed ***Hubel and Wiesel (1965)*** in focusing on how neurons aggregate inputs tuned to simpler features, the nonlinear mechanisms of spike generation, which determine how these inputs are summed, are also thought to be important ***(Riesenhuber and Poggio, 1999)***. The present study supports this idea by showing how a single voltage-gated current, ***I_K_LT__***, has the potential to dramatically shift how information about stimulus identity is encoded. As we become increasingly aware of the diversity of cell types in the brain ***(Wang et al., 2006; van Aerde and Feldmeyer, 2015; Tasic et al., 2016)*** and the activity-dependent mechanisms that can modulate intrinsic electrophysiological properties ***(Nataraj and Turrigiano, 2011; Mahon and Charpier, 2012; Dehorter et al., 2015)***, it is important to account for intrinsic dynamics in models of sensory processing.

## Methods and Materials

### Linear-Dynamical Cascade Model

#### Biophysical model

The model used in this study is a conductance-based, single-compartment model for CM neurons ***(Chen and Meliza, 2017)***, which was adapted from the ventral cochlear nucleus model of ***Rothman and Manis (2003)***. The currents in the model include four voltage-gated potassium and sodium currents, a leak current, and a hyperpolarization-activated cation current. A key component of the model is the low-threshold potassium current (***I_K_LT__***), which determines whether the neuron produces tonic or phasic responses to step depolarization. When (the maximal conductance of ***I_K_LT__***) is low, the model neuron produces sustained responses to weak and moderate depolarizations; when ***g_K_LT__*** is high, the model only fires phasically, at the onset of the current step. The model parameter values follow ***Rothman and Manis (2003)***, with a few adjustments to match the resting potential and spike threshold for CM neurons. The calculations presented here used the consensus model parameters from ***Chen and Meliza (2017)*** for tonic and phasic cells. The model simulation code was generated using spyks (https://github.com/melizalab/spyks; version 0.6.6), and the dynamics were integrated using a 5th-order Runge-Kutta algorithm with an adaptive error tolerance of 1 × 10^-5^ and an interpolated step size of 0.025 ms.

#### Auditory response simulation

To simulate an auditory response, the model was driven by a current, ***I***_*stim*_(*t*), which was calculated from the convolution of a spectrotemporal receptive field (RF)with the spectrogram of an auditory stimulus. In this formulation, the RF approximated the synaptic drive to the neuron as a weighted sum of linear, time-invariant filters over different frequency channels. To simulate stimulus-independent variability in the response, a second driving current, ***I***_*noise*_(*t*), was added. The noise current was randomly generated pink noise (1/*f* power spectrum), low-pass filtered at 100Hz and scaled relative to the signal to achieve a set signal-to-noise ratio (SNR). For all of the analyses presented in this paper with the exception of the analysis of the effect of the signal-to-noise ratio, SNR was set at 4.

RFs were constructed with a Gabor filter based on ***Woolley et al. (2009)***:

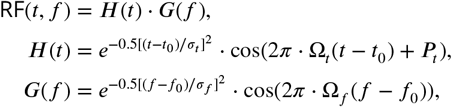

where ***H*** is the temporal dimension of the RF, ***G*** is the spectral dimension of the RF, *t*_0_ is the latency, *f*_0_ is the peak frequency, *σ_t_* and *σ_f_* are the temporal and spectral bandwidths, Ω_*t*_ and Ω_*f*_ are the temporal and spectral modulation frequencies, and ***P***_*t*_ is the temporal phase. Parameter values were randomly drawn from distributions set to match the modulation transfer function (MTF) of the RF ensemble to the MTF of zebra finch song ***(Singh and Theunissen, 2003; Woolley et al., 2009)***. The integral of each RF was normalized to one.

The models’ responses were simulated using 30 zebra finch songs recorded from our colony. Each recording was comprised of a single song motif repeated twice. Recordings were normalized to the same RMS amplitude and edited to be 2.025 s long, with 50 ms of microphone noise at the beginning to pad the convolution, and scaled to a consistent RMS amplitude. Start and end times of syllables were identified by visual inspection. Repeated syllables were grouped in the decoding analyses. Spectrograms of the stimuli were calculated using the short-time Fourier transform algorithm with a Hanning window of 256 points and then resampled to give a frequency resolution of 50 channels between 0 and 8 kHz. Successive windows were spaced 1.0 ms apart.

In the context of this simulation, a model neuron was defined as the combination of one RF and one biophysical parameter set (phasic or tonic). 60 RFs were generated to produce paired phasic and tonic simulations (*n* = 120 neurons or 60 pairs). For the analysis of the relationship between selectivity and temporal modulation, 8 RFs were generated and for each RF Ω_*t*_ was set to 10,20,…, 80 (*n* = 128 neurons or 8 sets of pairs). The 30 zebra finch songs were presented 10 times each to each neuron with different values of ***I***_*noise*_(*t*) producing trial-to-trial variability. Noise currents in each trial were identical between paired phasic and tonic neurons. The total amplitude of the convolution was normalized by the bandwidth of the RF on the frequency axis (*σ_f_*) to account for the differences in amplitudes between narrowband and broadband RFs. The output of the model was a simulated voltage trace.

#### Data analysis

Spike times were extracted from the simulated responses using a simple window discriminator (https://github.com/melizalab/quickspikes; version 1.3.3). We calculated spike rate, ***r***_*ij*_, as the number of spikes evoked by syllable *i* in trial *j*, divided by the duration of the syllable. Selectivity was quantified using activity fraction ***(Rolls and Tovee, 1995; Meliza and Margoliash, 2012)***, a nonparametric index defined as:

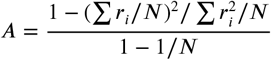

where ***r***_*i*_ is the rate for syllable *i* averaged across trials, and ***N*** is the total number of syllables.

Mutual information (MI), total entropy, and noise entropy were calculated following ***Jeanne et al. (2011)***. Response rates were discretized into 15 bins between the minimum and maximum rate of the model. Total entropy was calculated as ***H***(***R***) = −Σ*p*(***r***)log_2_*p*(***r***), noise entropy as ***H***(***R|S***) = −Σ*p*(*s*)Σ*p*(***r***|*s*) log_2_*p*(***r|s***), and mutual information as ***I***(***R; S***) = ***H***(***R***) – ***H***(***R|S***), where ***r*** is the rate and *s* is the syllable. Because of the large number of stimuli and trials, and because we were interested in differences between models presented with exactly the same stimuli, we did not correct entropy or MI for sample size bias.

#### Stimuli

Thirty male zebra finches provided song recordings that were used as stimuli in the simulation experiments. All animal use was performed in accordance with the Institutional Animal Care and Use Committee of the University of Virginia. Adult zebra finches were obtained from the University of Virginia breeding colony. During recording, zebra finches were housed in a soundproof auditory isolation box (Eckel Industries, Cambridge, MA) with *ad libitum* food and water and were kept on a 16:8h light:dark schedule. A mirror was added to the box to stimulate singing. Recordings were made with an Audio-Technica Pro 70 microphone, digitized with a Focusrite Scarlett 2i2 at 44.1 kHz, and stored to disk using custom C++ software (https://github.com/melizalab/jill; version 2.1.4). A typical recording session lasted 2-3 days. A single representative song was selected from each bird’s recorded corpus and was high-pass filtered at 500Hz with a 4th-order Butterworth filter.

#### Extracellular data

Analyses based on extracellular data were performed on the publicly available datasetfrom ***Theunissen etal. (2011)*** available at http://crcns.org/data-sets/aa/aa-2. Neural recordings were collected from adult male zebra finches as described in ***Gill et al. (2006)***. Only responses from CM neurons presented with conspecific song were used these analyses (*n* = 37). Selectivity and MI analyses were performed as described above, except that 10 response bins were used for MI analysis instead of 15, due to the smaller stimulus set.

## Acknowledgments

We thank Tyler Robbins for assistance in model development and JC Cang and Alev Erisir for critical comments and discussions.

